# NIR-II fluorescence microscopic imaging of cortical vasculature in non-human primates

**DOI:** 10.1101/853200

**Authors:** Zhaochong Cai, Liang Zhu, Mengqi Wang, Anna Wang Roe, Wang Xi, Jun Qian

## Abstract

Vasculature architecture in the brain can provide revealing information about mental and neurological function and disease. Fluorescence imaging in the second near-infrared (NIR-II) regime with less light scattering is a more promising method for detecting cortical vessels than traditional visible and NIR-I modes. Here, for the first time, we developed, NIR-II fluorescence microscopy capabilities for imaging brain vasculature in macaque monkey. The first is a wide-field microscope with high temporal resolution (25 frames/second) for measuring blood flow velocity and cardiac impulse period, and the second is a high spatial resolution (<10 μm) confocal microscope producing three-dimensional maps of the cortical microvascular network (∼500 μm deep). Both were designed with flexibility to image various cortical locations on the head. Use of a clinically approved dye provided high brightness in NIR-II region. This comprises an important advance towards studies of neurovascular coupling, stroke, and other diseases relevant to neurovascular health in humans.

## Introduction

Fluorescence bioimaging is a sensitive, non-invasive and radiation-free technology that is capable of revealing biological structure and function^1^. Its utility has been demonstrated in clinical applications, such as imaging-guided surgery for liver tumors and retinal angiography^2, 3^. However, its application for studying microvascular architecture and function in the human brain has not been well developed. There is now rapidly growing recognition that health of brain vasculature is essential for normal neurological and mental function. In particular, the role of fine microvasculature is proving to be a critical component in meeting the oxygen and energy demands of normal neuronal function^4–7^. Decline of vascular health can disrupt neurovascular coupling, leading to oxygen delivery mismatches and resulting functional decline. Indeed, common neurological disorders such as Alzheimer’s or ALS (amyotrophic lateral sclerosis) may be rooted in disorders of vascular structure and/or function^8, 9^. The purpose of this study is to develop a fluorescence microscopic imaging methodology for studying vascular architecture and function for future clinical application.

A traditional way to image vasculature is based on fluorescence signals whose wavelength is in the visible spectral range (400–760 nm) or first near-infrared (NIR-I) spectral region (760–900 nm)^10^. Both methods have small tissue penetration depth and low spatial resolution due to the severe scattering of fluorescence photons by biological tissues^11^. A more recent development, fluorescence bioimaging in the second near-infrared spectral region (NIR-II), takes advantage of emission wavelengths in the 900-1700 nm spectral range. This fluorescence bioimaging method utilizes long-wavelength and low-scattering signals, providing larger penetration depth and higher spatial resolution than visible or NIR-I fluorescence bioimaging^12^. Given that biological tissue usually has low autofluorescence in the long wavelength regime, NIR-II fluorescence bioimaging also provides high signal to background ratio (SBR)^13^.

These advantages have made NIR-II fluorescence bioimaging a method of choice for functional applications in mice, including whole-body angiography, organ visualization, as well as diagnosis and imaging-guided treatment of tumours^10–12, 14–22^. With respect to brain vasculature, NIR-II fluorescence microscopy has been used in studies of vascular structural architecture and blood flow velocity of mice^23^. Such studies in mice have been extremely exciting and have pushed forward the field. However, given the significant differences in structural and functional aspects of brain vasculature in mice vs humans, developing NIR-II fluorescence microscopy for human clinical use still requires development.

Here, to enable future clinical applications, we have developed NIR-II methods for brain micro-angiography in the rhesus macaque monkey, an animal model with cortical structure and organization similar to humans. One significant difference between primates (both human and nonhuman primates) and mice is that much of the cerebral cortex is organized into modular functional units of integrated neuronal response (termed columns or domains)^24–26^. This primate-specific functional architecture suggests that there may be specializations of vascular architecture to meet the metabolic and hemodynamic demands of columnar unit-based response, a possibility that is receiving growing attention^27–30^. As a step towards clinical application, our goal was thus to develop a method appropriate for the nonhuman primate, one that meets the needs of studying vascular structure and function. We aimed to achieve high spatial resolution for observing even the smallest vessels (5-10 μm), high temporal resolution for monitoring blood flow (up to 0.6 mm/s) in capillaries, and depth of imaging permitting visualization of the superficial layers of the cerebral cortex. Moreover, we intend this method to be relatively easy to use and at moderate cost, important for developing a practical and robust tool for clinical use.

Here, we report the development of two NIR-II fluorescence microscopes adapted to brain imaging of large animals. Using a clinically approved dye named indocyanine green (ICG) as a bright NIR-II fluorescent probe^31, 32^ in rhesus macaques, we demonstrate that the wide-field NIR-II fluorescence microscope offers high temporal resolution^23^, sufficient for realizing real-time imaging of cerebral blood vessels and achieving measurement of CBF velocity and cardiac cycle. The second microscope, a confocal NIR-II fluorescence microscope, offers high spatial resolution and high SBR (signal to background ratio) via optical sectioning capability^33^, enabling reconstruction of a clear three-dimensional volume of cortical vasculature up to an imaging depth of ∼500 μm. Capillary vessels with a diameter less than 7 μm could be resolved.

## Results

### Optical characterization of ICG in rhesus macaque serum

Key to clinical feasibility is the availability and applicability of safe and effective labelling agents. Thus far, several types of NIR-II fluorescent probes, including carbon nanotubes^34^, rare-earth doped nanoparticles^35^, quantum dots^36^, small-molecule organic dyes^37^ and organic nanoparticles^38, 39^, have been utilized for *in vivo* imaging of small animals. However, these probes have not been developed for human use. To work towards a capability that is easily compatible with human use, we utilized indocyanine green (ICG, Fig. 1a), a probe that is readily available and is approved by US Food and Drug Administration (FDA). ICG has been previously used in humans as a typical NIR-I fluorophore (for studies of liver and retina)^2, 3, 40^, and, as been previously shown, also offers the advantage of bright NIR-II fluorescence emission^31^.

**Fig. 1.**
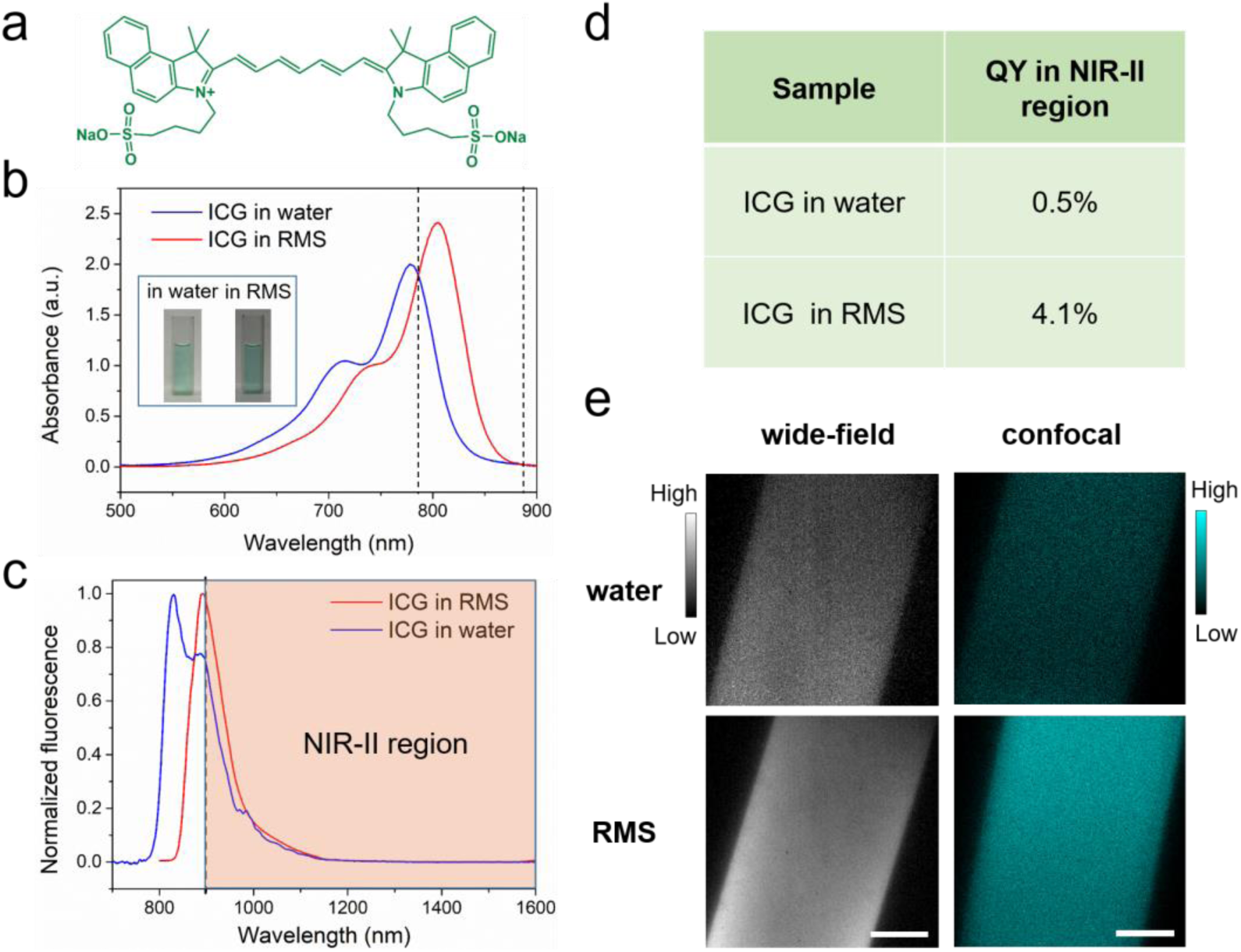
Optical characterization of ICG in RMS. **a** Molecular structure of ICG. **b** The absorption spectra of ICG in water (blue) and RMS (red). Inset: bright-field images of ICG in water and RMS. Dashed lines: range of increased absorbance in RMS. **c** Normalized fluorescence spectra of ICG in water and RMS. Shaded region: fluorescence spectrum of ICG in RMS beyond 900 nm. **d** NIR-II fluorescence quantum yield (QY) of ICG in water and ICG in RMS. **e** Images of ICG in water and RMS from NIR-II fluorescence wide-field microscope (left panels, 780 nm LED excitation) and confocal microscope (right panels, 793 nm laser excitation). Scale bars: 100 μm.

Prior to *in vivo* applications in imaging cortical vasculature of rhesus macaques, we first evaluated the optical characteristics of ICG in rhesus macaque serum (RMS). We found two important benefits of ICG. First, by comparing the absorption spectra of ICG in water with that in RMS (Fig. 1b), we observed that the absorption of ICG in RMS peaked at 806 nm, a wavelength that is red-shifted ∼30 nm compared to that of ICG in water. This enhanced absorbance in RMS (in the range of 786-888 nm) significantly increases the excitation efficiency of 793 nm laser/780 nm LED, important for the feasibility of following *in vivo* imaging in primates. Second, we found that the normalized fluorescence spectra (Fig. 1c) of ICG in RMS showed an emission peak at ∼900 nm, which is ∼60 nm red-shifted compared to ICG in water. This increases greatly the portion of fluorescence emission in NIR-II spectral region (shaded region of graph). Moreover, a NIR-II fluorescence quantum yield (QY) of ICG in RMS reaches 4.1%, which is a dramatic increase of 8.2-fold compared to ICG in water (Fig. 1d); this is also higher than most other NIR-II fluorescent probes^34, 35, 37, 41^. The increased absorbance and QY result in striking improvements in signal brightness as seen through the NIR-II fluorescence wide-field microscope (Fig 1e, left panels, 98% increase) and confocal microscopic imaging results (Fig. 1e, right panels, 214% increase). The unique optical feature of ICG in RMS is due to the fact that ICG molecules (molecular size less than 0.5 nm) tend to adsorb on serum proteins and form an “organic dye-protein complex” (Supplementary Fig. 1), which thus reduces the aggregation-caused quenching (ACQ) effect of ICG arising from its self-aggregation in water^22^.

### *In vivo* NIR-II fluorescence wide-field microscopic imaging of cerebral blood vessels of rhesus macaques

NIR-II fluorescence wide-field microscopy offers high temporal resolution and easy operation, which are features of traditional wide-field fluorescence microscopy. In addition, due to the minimal scattering of fluorescence signals in biological tissues, NIR-II fluorescence wide-field microscopy achieves larger penetration depth. Though there have been several reports on *in vivo* NIR-II fluorescence wide-field microscopic imaging in mice (e.g. brain and tumour angiography)^23, 41, 42^, such imaging on large animals has never been demonstrated. To establish such a capability, we custom-designed a NIR-II fluorescence wide-field microscopic system.

The configuration of our lab-built optical setup is shown in Fig. 2a. A 780 nm LED was coupled to a microscope illuminator and utilized as the excitation source. NIR-II fluorescence images were collected by an NIR anti-reflection objective and recorded with an InGaAs camera. One challenge for imaging large animal brains is that, unlike mice, their cranial windows are not always horizontal. Due to the large brain size, areas of interest may lie in planes of antero-posterior or mediolateral tilt. To address this need, we mounted the whole optical system on a multi-direction adjustable shelf (yellow shaded area in Fig. 2a), thereby enabling convenient translation and rotation, as demonstrated by the straight (green) and curved (blue) arrows. This permitted positioning of the objective directly above and perpendicular to the cranial window. In addition, for micropositioning of the field of view during imaging (red arrows), the rhesus macaque was placed on a stage which could be moved in the x-y plane. For fine z-direction adjustment, the objective was fixed on a motor-driven electric module capable of a 10 μm step-by-step tomographic imaging. This instrument was also designed with the potential of imaging in awake, behaving primates with head-fixation.

**Fig. 2.**
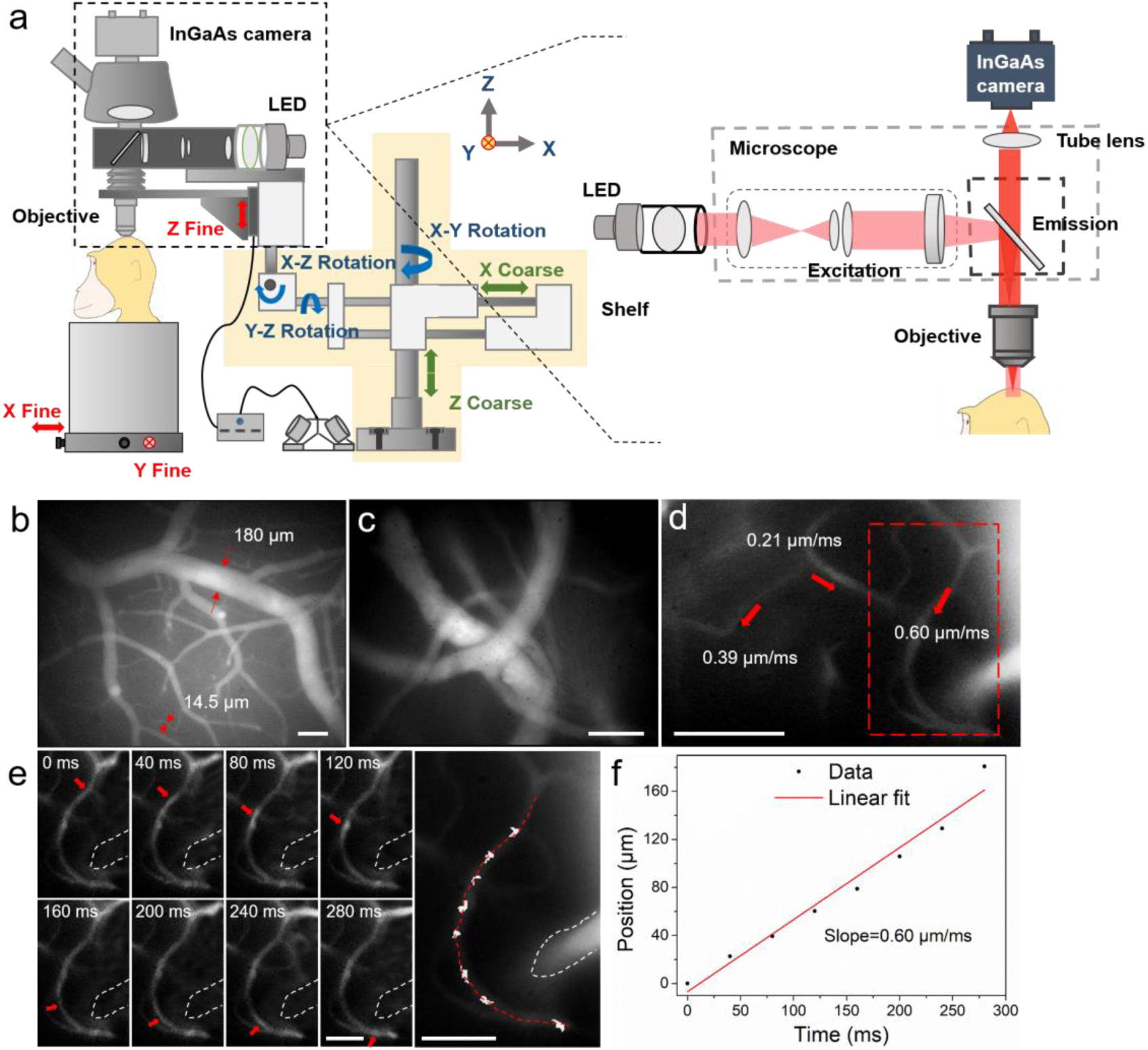
*In vivo* fluorescence wide-field microscopic imaging of cerebral blood vessels of the rhesus macaque in the NIR-II spectral region. **a** Schematic illustration of the custom NIR-II fluorescence wide-field microscopic imaging system. Red arrows: fine adjustment. Blue arrows: rotation. Green arrows: coarse adjustment. The excitation and imaging light paths are showed on the right panel. **b** A typical microscopic image of brain blood vessels with low magnification (∼3×). **c** A typical microscopic image of brain blood vessels with high magnification (∼25×). **d** Blood flow velocities in three sampled vessels. Velocity calculation shown in **e**, **f**. Red arrows indicate the directions of blood flow. Dashed red box: region shown in **e**. **e** Frames showing tracking a fluorescent point in a capillary (diameter = 8 μm). The figure on the right marks the tracked positions. **f** A plot of the position of the point as a function of time. The linear fit (r =0.986) reveals an average blood velocity of 0.60 μm/ms in the capillary. Excitation wavelength: 780 nm. Exposure time of the InGaAs camera: 10 ms. Scale bars in **b**, **c** and **d**: 100 μm, scale bars in **e**: 50 μm.

Following implantation of a cranial window under anesthesia (see Methods), ICG was intravenously injected (1.43 mg/kg, below clinical safe dose of 5 mg/kg^43^) and the brain was imaged through the custom NIR-II fluorescence microscope. A low-magnification objective (∼3×) was first used to provide a structural map of cerebral blood vessels (Fig. 2b). Due to the large field of view (FOV, 2.1×1.7 mm) and bright NIR-II fluorescence signals from ICG, both large blood vessels (diameter=180 μm) and microcapillaries (diameter=14.5 μm) could be visualized simultaneously. This map was then used to select and focus on a specific sub-region of interest (ROI) via a ∼25× objective for obtaining a higher-resolution image of blood vessels (Fig. 2c, depth= 130 μm).

Our next goal was to understand functional aspects of the vascular network by studying the flow within different sized vessels. Under the ∼25× objective, numerous bright points from NIR-II fluorescence emission of ICG were observed moving along capillaries (Fig. 2d and Supplementary MOV S1). Under baseline conditions (e.g. under anesthesia), flow velocity within single microvessels should be stable. To calculate CBF velocity, a single point on a capillary was selected (Fig. 2d, red dashed box) and continuously tracked at 25 frames per second (FPS) (Fig. 2e). As shown in Fig. 2f, by plotting the positions of this point at different frames and performing a linear fit, we determined a CBF velocity of 0.60 μm/ms for this fine vessel. The blood flow velocities in other two capillaries within the FOV were also determined in the same way (0.21 μm/ms and 0.39 μm/ms, red arrows in Fig. 2d). In this way, the flow characteristics of multiple capillaries in a field of view could be evaluated at the same time. Since veins collect blood from branches while arteries distribute blood into branches^28^, blood vessel type (vein or artery) could be recognized according to the flow direction (Supplementary Fig. 2 and Supplementary MOV S2).

In the z direction, blood vessels at various depths were imaged (Supplementary Fig. 3). The images at two typical depths were analyzed and the results illustrated the microcapillaries could be resolved (FWHM=7.8 μm at the depth of 300 μm and FWHM=8.5 μm at the depth of 170 μm, Supplementary Fig. 4). Nevertheless, since “wide-field excitation and area detection” imaging mode was adopted, the fluorescence signals both above and below the focal plane of objective were also recorded and became the background of the image on the focal plane, which reduced the SBR of imaging.

In addition, in some large blood vessels, regions with bright/dark borders were clearly observed oscillating in position over time with cardiac impulse (Supplementary MOV S3 clearly shows this border pulsation). Supplementary Fig. 5a illustrates the highest (top panel) and the lowest (bottom panel) position of the bright/dark border during one cardiac impulse period. The fluctuation of the bright/dark border position is displayed in Supplementary Fig. 5b. By plotting the timepoints of peaks of each pulse and making a linear fit, a pulse rate of 2 Hz was obtained (Supplementary Fig. 5c). This calculated period was consistent with that concurrently measured by the physiological monitor (120 pulses per minute). This demonstrates that the high temporal resolution of NIR-II fluorescence wide-field microscopy provides simultaneous pulse rate information from large vessels while monitoring blood flow velocity in small vessels; these are parameters which are likely to prove useful in studies of vascular flow and hemodynamics.

### *In vivo* NIR-II fluorescence confocal microscopic imaging of cerebral blood vessels of rhesus macaques

Although our NIR-II fluorescence wide-field microscopy was capable of realizing real-time brain imaging on rhesus macaques, its spatial resolution and SBR were inevitably influenced by out-of-focus signals (fluorescence signals both above and below the focal plane of objective). To address this challenge, we turned to NIR-II fluorescence confocal microscopy, which offers fine optical sectioning and high SBR, as well as large penetration depth from NIR-II fluorescence bioimaging, as demonstrated by *ex vivo* and *in vivo* bioimaging in mice^17, 36, 44, 45^. To achieve this capability in large animals, we custom-designed a NIR-II fluorescence confocal microscopic system, modified from our previous lab-built setup^33, 41^. As shown in Fig. 3a, to achieve point detection of NIR-II fluorescence signals, we used a 793 nm laser as the excitation source and an InGaAs photomultiplier tube (PMT) with high QY at 900-1700 nm wavelengths. A multimode fiber with a core diameter of 400 μm played the role of a pinhole to exclude the out-of-focus signals (signals produced outside the focus of excitation laser). The galvanometers inside the scan unit could achieve fine point-by-point scanning in the x-y direction. The vertical position of the objective was adjusted by a motor-driven electric module to realize fine step-by-step (10 μm) z-direction movement. The whole confocal microscopic system was fixed on a multi-direction adjustable shelf, which permitted accurate positioning via convenient translation and rotation of the objective directly above and perpendicular to the cranial window (red arrows in Fig. 3a). The subject was placed on a stage which could be moved in the x-y direction prior to imaging.

**Fig. 3.**
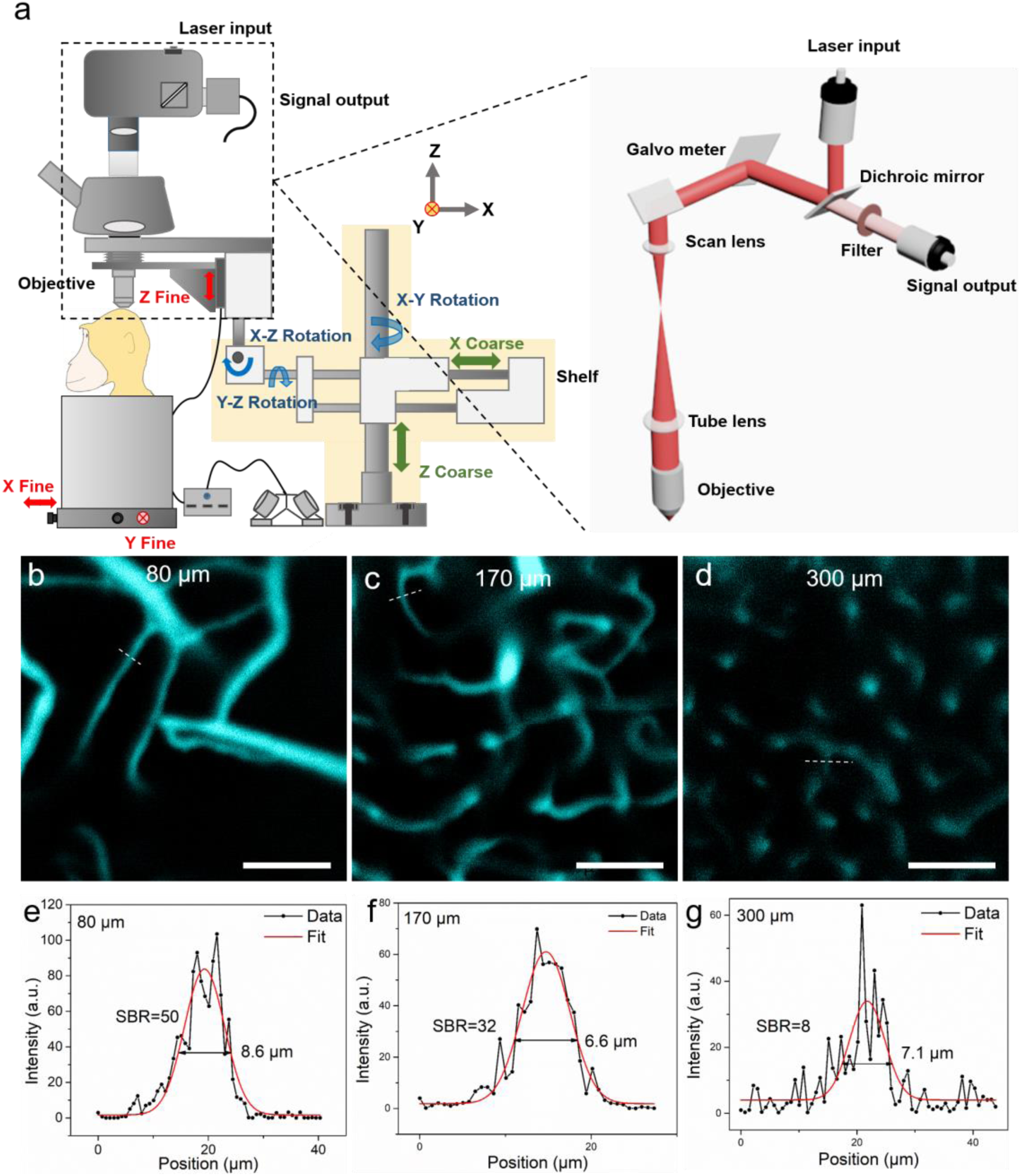
NIR-II fluorescence confocal microscopic *in vivo* imaging of cerebral blood vessels of the rhesus macaque with high spatial resolution and SBR. **a** The schematic illustration of the NIR-II fluorescence confocal microscopic imaging system. Red arrows: fine adjustment. Blue arrows: rotation. Green arrows: coarse adjustment. The excitation and imaging light paths are showed on the right panel. **b-d** NIR-II fluorescence confocal microscopic images at three typical depths (80 μm, 170 μm and 300 μm). White dashed lines: locations of cross-sections depicted in **e-g**. **e-g** The cross-sectional fluorescence intensity profiles (black) and the related Gaussian fits (red) along the capillary vessels, taken from locations (white-dashed lines) in **b-d**. Excitation wavelength: 793 nm. Laser power: ∼40 mW before the objective. PMT voltage: ∼531 V. Pinhole diameter: 400 μm. All scale bars: 100 μm.

After testing the performance of this setup in mice (Supplementary Figures 6 and 7), *in vivo* cerebral cortical vessel imaging on the rhesus macaques was implemented (injected dose of ICG was 2.86 mg/kg, below clinical safe dose of 5 mg/kg). Due to the bright NIR-II fluorescence emission of ICG in RMS and the spatial filtering capability of NIR-II fluorescence confocal microscope, we achieved high spatial resolution and high SBR of the cortical vascular network (Fig. 3). Fig. 3b∼d are representative confocal images taken at depth of 80 μm, 170 μm and 300 μm, respectively, showing clear vascular morphology. In Fig. 3e∼g, profiles and Gaussian fits of fluorescence intensity profiles along the white dashed lines in Fig. 3b∼d are shown. At a depth of 170 μm (Fig 3c), the SBR was as high as 32, permitting vessels as small as 6.6 μm in diameter (FWHM) to be resolved. Even at a depth of 300 μm, the SBR of 8 was achieved, providing clear images of a capillary vessel with a diameter (FWHM) of 7.1 μm. Thus, in comparison to NIR-II fluorescence wide-field microscopy (Supplementary Fig. 4), the greatly improved SBRs (2.1 at the depth of 170 μm and 1.5 at the depth of 300 μm) produced spatially resolved images of capillaries at good penetration depths.

Tomographic imaging of cerebral blood vessels was further conducted with a step resolution of 10 μm. The reconstructed images at various depths (0-130 μm, 140-250 μm and 260-470 μm) are shown in Fig. 4a∼c, where the cerebral blood vessels of rhesus macaque showed clear and distinct layers. By reconstructing the images, a vivid 3D vascular architecture was obtained (Fig. 4d, Fig. 4e and Supplementary MOV S4). A maximal imaging depth of 470 μm could still be 15 obtained, thanks to the “NIR-I excitation and NIR-II emission” imaging mode. In comparison to the NIR-II fluorescence wide-field microscopy, NIR-II fluorescence confocal microscopy possessed higher SBR and spatial resolution, though at the price of lower temporal resolution (5.24 s per picture). Despite this, a scanning speed of 20 μs/pixel was achieved, a speed still much faster than that (50 μs/pixel) achieved in other studies in mice^17^. We conclude that our NIR-II fluorescence confocal microscope is capable of high spatial resolution 3D imaging of vascular networks in rhesus macaques.

**Fig. 4.**
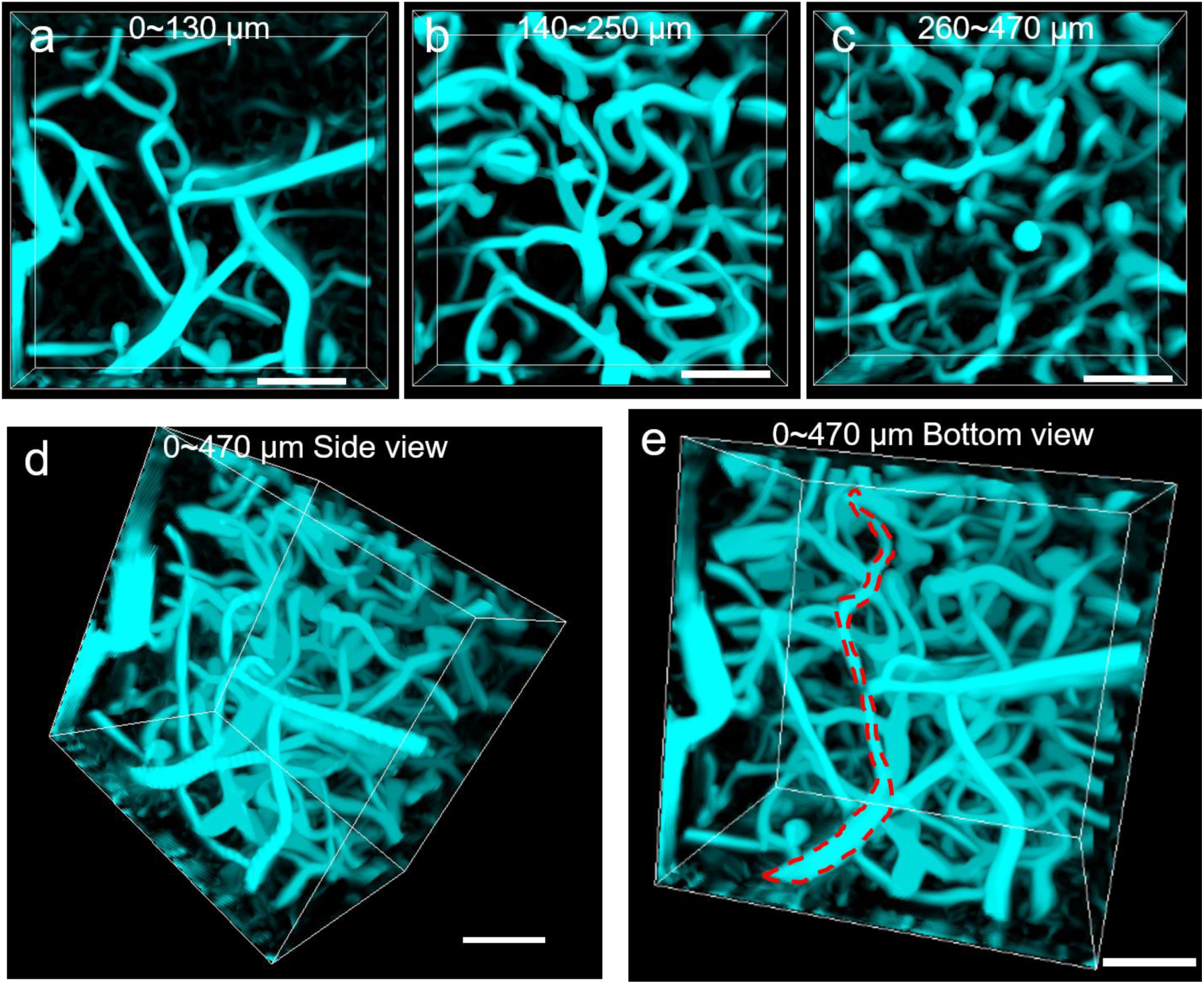
NIR-II fluorescence confocal microscopic *in vivo* imaging of cerebral blood vessels of the rhesus macaque with large penetration depth. **(a-c)** 3D reconstructed NIR-II fluorescence confocal microscopic images of brain blood vessels at various depths (**a**: 0-130 μm, **b**: 140-250 μm, **c**: 260-470 μm). (**d-e**) 3D reconstructed NIR-II fluorescence confocal microscopic images up to depth of 470 μm from the side view **d** and bottom view **e**. Red dashed lines in **e**: outline of single vessel. Excitation wavelength: 793 nm. Laser power: ∼40 mW before the objective. PMT voltage: ∼531 V. Pinhole diameter: 400 μm. All scale bars: 100 μm.

## Discussion

### A new method for high spatial and temporal resolution angiography in primates

In this study, we provide a promising NIR-II fluorescent approach for cortical angiography in nonhuman primates. For the first time, we conducted *in vivo* NIR-II fluorescence bioimaging of cerebral cortical microvasulature in rhesus macaques. To achieve structural and functional imaging on cerebral vascular network, we specifically designed two NIR-II fluorescence microscopes. Both of them adopted NIR-I light source to reduce the tissue-absorption of excitation, and used the InGaAs detector to collect NIR-II signals with a minimum of tissue-scattering, thereby enabling large imaging depth. The NIR-II fluorescence wide-field microscope adopted the “area-excitation and area-detection” approach, which offered advantages of high temporal resolution and easy operation. The NIR-II fluorescence confocal microscope was based on “point-excitation, point-detection and spatial filtering”, with advantages of fine optical sectioning and high SBR.

In addition, we designed the microscopes so that they could image the brain at different angles, to permit study through cranial windows in different cortical areas of interest in rhesus macaques. Such flexibility will also prove valuable for future studies in awake, behaving monkeys. To overcome these difficulties, both microscopes were fixed on a multi-direction adjustable shelf, where translation and rotation were enabled and the objective angle and position could be flexibly adjusted above and perpendicular to the cranial window. In addition, the rhesus macaque was placed on a large and stable stage, where precise x-y movement was enabled. As far as we know, these are the first NIR-II fluorescence microscopes for *in vivo* imaging in non-human primates.

### Probe that is clinically compatible and bright

In addition to the development of the NIR-II fluorescence microscopic systems, NIR-II fluorescent probes were also critical for successful imaging in non-human primates. These probes must satisfy at least three requirements. The first is their bio-safety. Similar to humans, the experimental ethics must be carefully considered. ICG, which was approved by FDA in 1958^46^, has been safely utilized in various clinical applications, and should also have negligible toxicity in rhesus macaques. Thus, the use of biocompatible fluorescent probes are an essential feature and test of this method. A second requirement is that the probe must have sufficiently high brightness in NIR-II spectral region. This is important for obtaining higher temporal and spatial resolution, as well as reducing the excitation intensity and other parameters (e.g. imaging time) of the imaging instruments. Finally, to permit application of the needed volumes in large animals (such as rhesus macaques which weigh 5-7 kg), the fluorescent probes should be easily accessible and not too costly^47^. The medical ICG we adopted was commercially available (in hospitals), more readily available than all the other reported NIR-II fluorescent probes, and moderate in cost. Thus, unlike carbon nanotubes, rare-earth doped nanoparticles, quantum dots, small-molecule organic dyes, and organic nanoparticles, these requirements are readily met by ICG.

We further learned that ICG is optically advantageous for use in nonhuman primates. As a typical kind of NIR-I fluorescent probe^40^, ICG has been proven to exhibit bright NIR-II fluorescence^31, 48^, and it has extremely large absorbance and has been successfully employed for NIR-II fluorescence functional bioimaging in mice. We determined that the absorbance of ICG in RMS increased in the spectral region between 786 nm and 888 nm, compared to that of ICG in water. In addition, we measured a dramatic 8.2-fold increase of the NIR-II fluorescence QY of ICG in RMS compared to that of ICG in water. Thus, under the excitation of 780 nm LED (for wide-field microscope) or 793 nm laser (for confocal microscope), ICG in RMS exhibited much brighter NIR-II emission, making it ideal for *in vivo* cerebral blood vessel imaging of nonhuman primates.

### Implications and significance

We believe that there are important implications of a primate-compatible method for imaging fine vascular networks. The role of fine vasculature is central to both normal brain function and brain disease. For example, the fine capillary bed and associated precapillary arterioles are believed to be the center of essential oxygen delivery to cortical neurons^28^. Normal hemodynamic function may be dependent on distribution of vascular control points and key temporal control of local constriction and dilation in response to metabolic demand^4, 5^. Unlike mice, in primates and humans, many cortical areas are organized in so-called cortical columns, which are units of coherent neuronal function^24–26^. However, what the organizing principle of this coherence is still under investigation. The coherence of neuronal response suggests that the column may be a unit of oxygen demand, one which is potentially related to the modular architecture of vascular networks. In addition, as feedforward vs feedback cortical processes central to cognitive control are organized in differential laminar fashion^49^, understanding the laminar organization of microvasculature may be key to the cortical processes^50^. Thus, it is critical to have a method for studying architectural and functional aspects of the microvascular network in both the columnar and laminar dimensions in nonhuman primates. This capability will advance our understanding of human neurovascular function in normal brain function and in neurological and mental disease.

## Methods

### Materials

Indocyanine green was purchased from Dandong Yichuang Pharmaceutical Co. Ltd. China. Phosphate-buffered saline (PBS) was obtained from Sinopharm Chemical Reagent Co. Ltd. China.

### Optical characterization of ICG

Absorption spectra of ICG were obtained with a UV–vis–NIR scanning spectrophotometer (UV2550, Shimadzu, Japan). Fluorescence spectra were measured by a NIR spectrometer (FLS980, Edinburgh instruments, UK). Absolute NIR-II fluorescence quantum yield was measured by a QY measuring instrument (Quantaurus-QY C13534, Hamamatsu, Japan).

### Animals

All experimental procedures conducted in mice and monkeys were approved by the Institutional Animal Care and Use Committee of Zhejiang University and in accordance with the National Institutes of Health Guidelines. Male C57BL6/J mice (Jackson Labs; 9-10 weeks old) were housed at 24°C with a normal 12 h light/dark cycle, and fed with water and food *ad libitum*. In our study, three healthy adult male rhesus macaques (4-5 years old and weighing 5-7 kg) were used, and they were housed at the Nonhuman Primate Facility at Zhejiang University.

### Monkey surgical procedure

Rhesus macaques were anesthetized with ketamine hydrochloride (10 mg/kg)/atropine (0.03 mg/kg) and maintained with propofol (5 mg/kg per hour iv; induction, 5 mg/kg) or sufentanil (2 to 4 μg/kg per hour iv; induction, 3 μg/kg) supplemented with isoflurane (0.5 to 2%). Animals were intubated and artificially ventilated. End-tidal CO_2_, respiration rate, SpO_2_, heart rate, electrocardiogram and rectal temperature were continuously monitored and maintained. The end-tidal CO_2_ was kept around 4% by adjusting the rate and volume of the ventilator, and the rectal temperature of 37.5°C to 38.5°C was maintained by an animal warming system. To access the cerebral cortex, a custom-made imaging chamber, with a transparent quartz glass was implanted over the cortex and secured by ceramic screws. This provided sufficient cortical stabilization for imaging. Following head fixation and prior to NIR-II fluorescence imaging, ICG was injected intravenously through the saphenous vein (∼2 mg/kg) to label the vasculature. Following completion of surgical and experimental procedures, monkeys were recovered and provided analgesics.

### NIR-II fluorescence wide-field microscopic imaging

A lab-built optical system was established to conduct NIR-II fluorescence wide-field microscopic brain imaging on the rhesus macaques. A 780 nm LED (M780L3-C1, Thorlabs, USA) was used as the excitation source. Reflected by a 900 nm long-pass (LP) dichroic mirror (DMLP900R, Thorlabs) and passing through an air objective lens (LSM03, WD = 25.1 mm, Thorlabs) or an infrared anti-reflection water-immersed objective lens (XLPLN25XWMP2, 25×, NA = 1.05, Olympus, Japan), the 780 nm beam illuminated onto the brain of the rhesus macaque. The excited NIR-II fluorescence was collected by the same objective lens, passing through the same 900 nm LP dichroic mirror and a 900 nm LP filter (FELH0900, Thorlabs), and finally was recorded by an InGaAs camera (SW640, Tekwin, China) via a built-in tube lens in a triocular (BX51, Olympus). The objective was fixed on a motor-driven electric module (ZFM2020, Thorlabs) and could be moved in the Z axis to focus at different depths of the brain. The whole microscope was fixed on a multi-direction adjustable shelf, which permitted careful positioning of the objective right above the window. A flexible translation and rotation system permitted precise position of the microscope perpendicular to the cranial window. In addition, placement of the rhesus macaque on an independently controlled stage provided precise x-y movement when conducting imaging.

### NIR-II fluorescence confocal microscopic imaging

A lab-built optical system was set to perform NIR-II fluorescence confocal microscopic brain imaging on the rhesus macaques. A 793 nm CW laser (Rugkuta Optoelectronics, China) beam was introduced into a scan unit (Thorlabs) and then reflected by a 900 nm LP dichroic mirror (DMLP900R, Thorlabs). After passing through the galvanometers, scan lens and tube lens, the excitation light was focused on the cranial window of rhesus macaque by an infrared anti-reflection water-immersed objective lens (XLPLN25XWMP2, 25×, NA = 1.05, Olympus). The emitting NIR-II fluorescence passed the aforementioned light path (from the 900 nm LP dichroic mirror to the objective) in counter direction, as well as a 900 nm LP filter (FELH0900, Thorlabs), and was finally directed into a NIR sensitive InGaAs photomultiplier tube (PMT, H12397-75, Hamamatsu) through a multimode fiber. The pinhole size depended on the aperture (core diameter) of this fiber, which, in this case was 400 μm. The output electric signal from the PMT was amplified with an amplifier (C12319, Hamamatsu) and acquired images reconstructed. Similar to the NIR-II fluorescence wide-field microscope, the objective was fixed on the motor-driven electric module (ZFM2020, Thorlabs) and the laser excitation beam could be focused at different depths of the cortex. The whole microscope was also fixed on the multi-direction adjustable shelf, enabling precise adjustment of the position between objective and the cranial window of the rhesus macaque. Animals (rhesus macaques or mice) were placed on an independently controlled stage, and fine point-by-point scanning in x-y direction was achieved via galvanometers.

### Image analysis

All data analyses were performed using MATLAB (R2018b; MathWorks) and IMARIS (Oxford Instruments). For vessels 3D reconstruction, images were processed with Simple Non Local Means (NLM) Filter (Christian Desrosiers, https://ww2.mathworks.cn/matlabcentral/fileexchange/52018-simple-non-local-means-nlm-filter) to reduce noise, and were enhanced structures in 2D images using hessian eigenvalues by Jerman Enhancement Filter (Tim Jerman, https://ww2.mathworks.cn/matlabcentral/fileexchange/63171-jerman-enhancement-filter). For processed images, IMARIS 3D Surfaces were used to precisely rebuild the 3D network map of blood vessels. In order to track the movements of bright points in the capillaries, the original images were subtracted to the average image of all the frames and the Matlab image tool box was used to obtain the bright points positions in each frames.

## Supporting information

MOV S1

MOV S2

MOV S3

MOV S4

## Acknowledgements

This work was supported by National Natural Science Foundation of China (61735016, 81430010, 61975172, 81702508, 91632105, 31627802) and Zhejiang Provincial Natural Science Foundation of China (LR17F050001, LY17C090005).

## Author contributions

J.Q., W.X., A.W.R., Z.C., L.Z. and M.W. conceived and designed the experiments. Z.C., L.Z. and M.W. performed the experients. J.Q., W.X., A.W.R., Z.C., L.Z. and M.W. analyzed the data. J.Q., W.X., A.W.R., Z.C. and L.Z. wrote the manuscript. All the authors contributed the discussions of the manuscript.

---

**Supplementary Figure 1.**
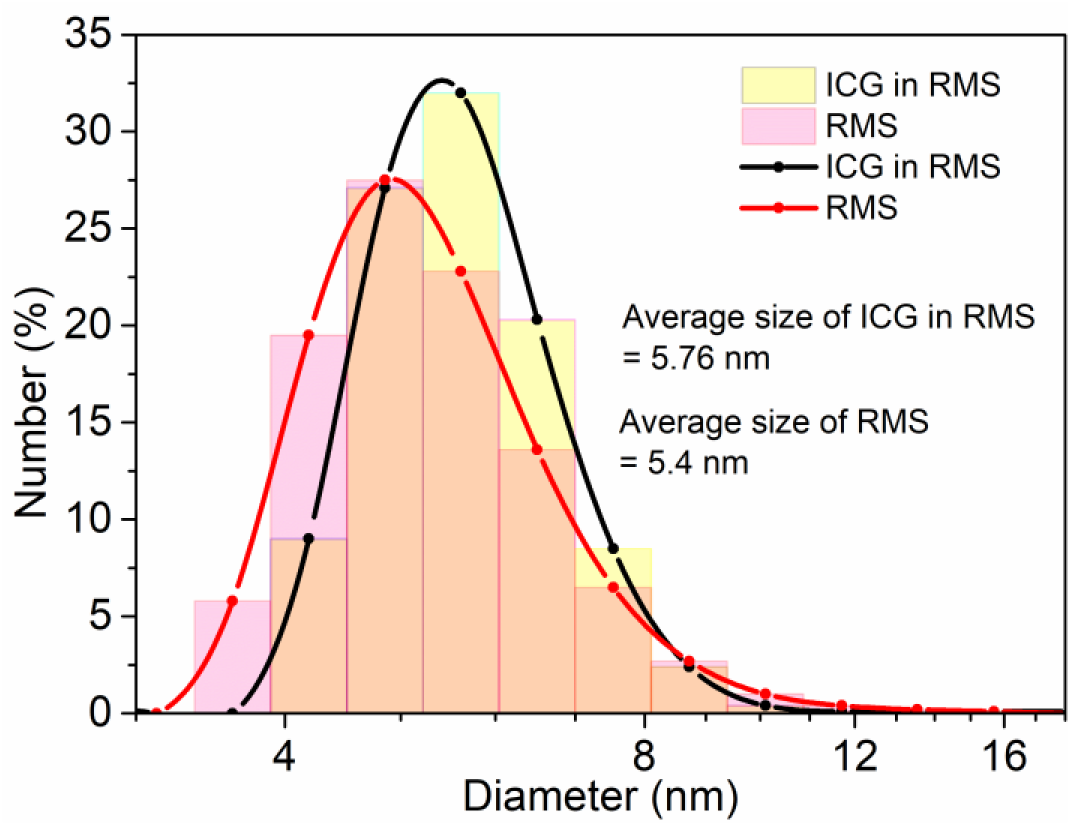
Hydrodynamic mean size distributions of rhesus macaque serum (pink) and ICG in rhesus macaque serum (yellow) by dynamic light scattering. These distributions are significantly different (RMS mean: 5.4 nm, ICG in RMS mean: 5.76 nm; (Χ^2^ = 131.6, p < 0.001).

**Supplementary Figure 2.**
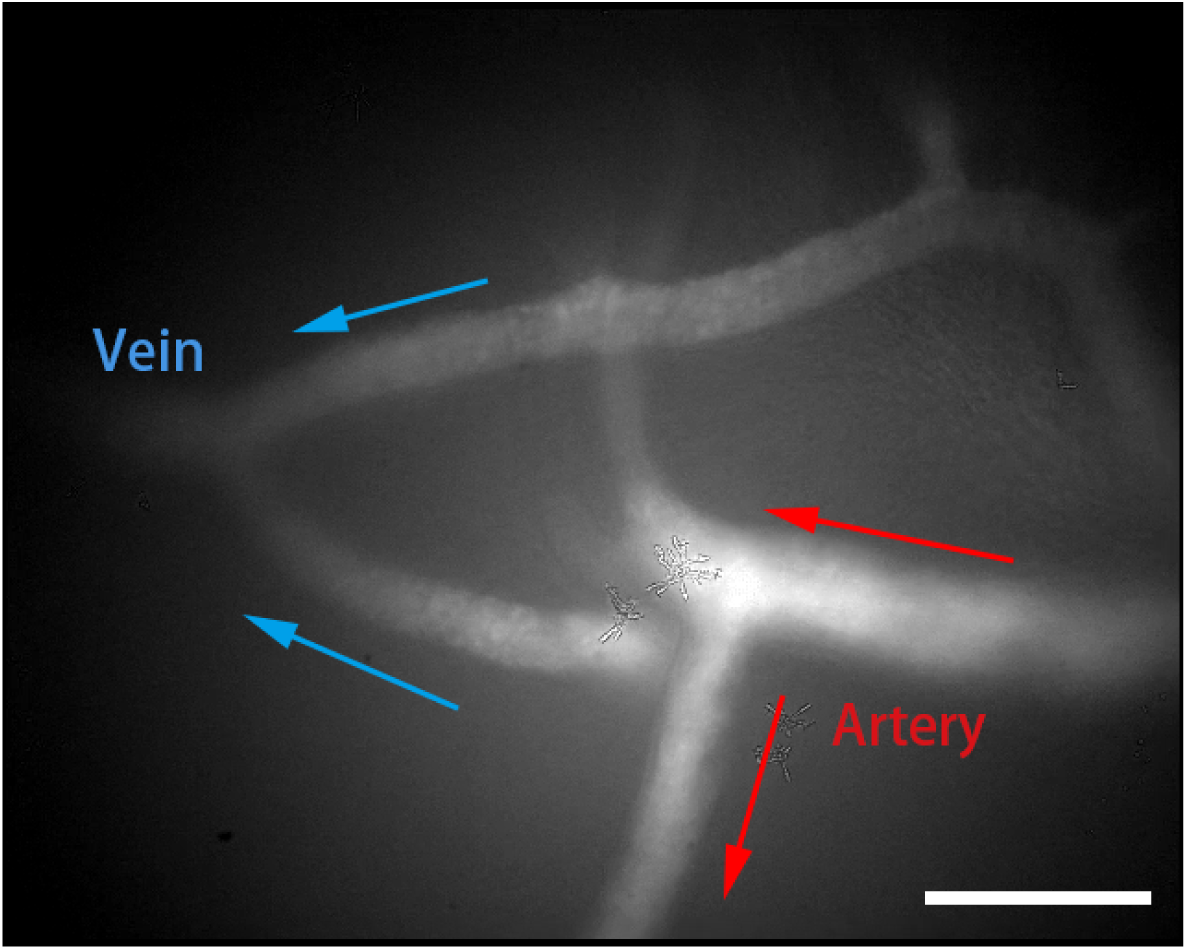
NIR-II fluorescence wide-field microscopic image of cerebral vessels showing the blood flow directions and determination of arteries and veins. Scale bar: 100 μm.

**Supplementary Figure 3.**
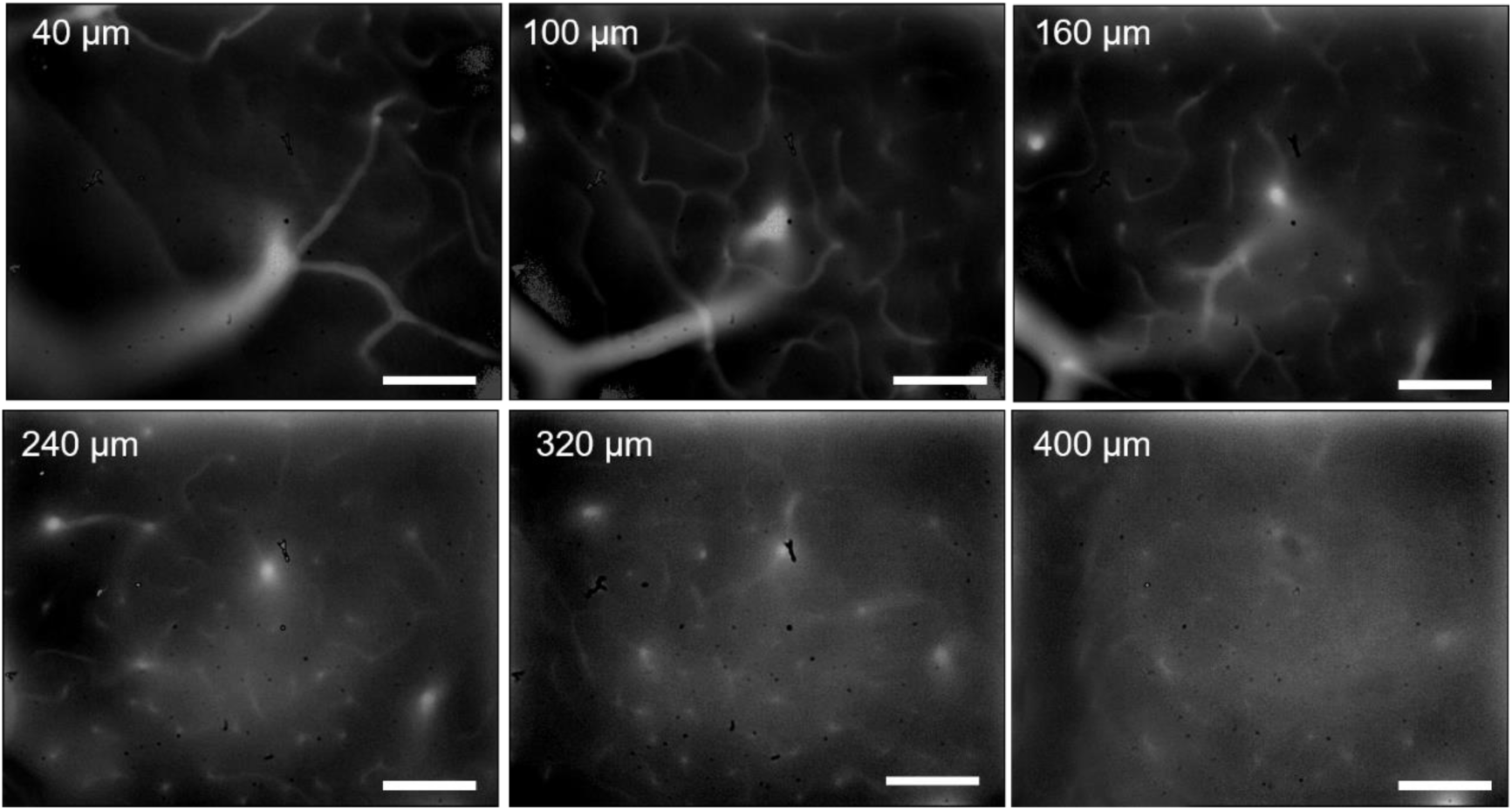
NIR-II fluorescence wide-field microscopic images of cerebral blood vessels of the rhesus macaque at various depths (40 μm, 100 μm, 160 μm, 240 μm, 320 μm, and 400 μm). Scale bars: 100 μm.

**Supplementary Figure 4.**
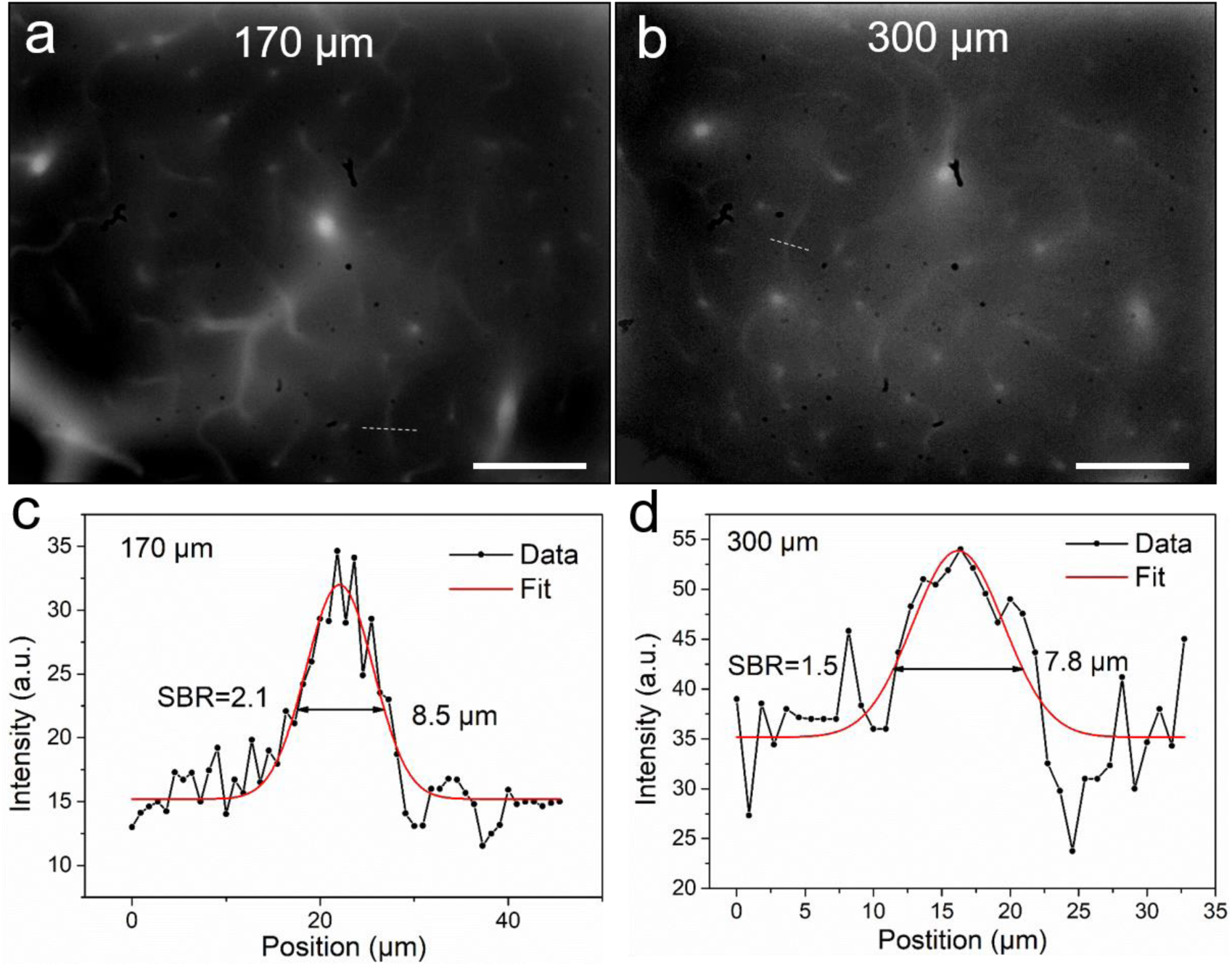
**a** and **b** NIR-II fluorescence wide-field microscopic images of cerebral blood vessels of the rhesus macaque at two typical depths (170 μm and 300 μm). **c-d** The cross-sectional fluorescence intensity profiles (black) and the related Gaussian fitting (red) along the capillaries indicated by the white-dashed lines in **a** and **b**. Scale bars: 100 μm.

**Supplementary Figure 5.**
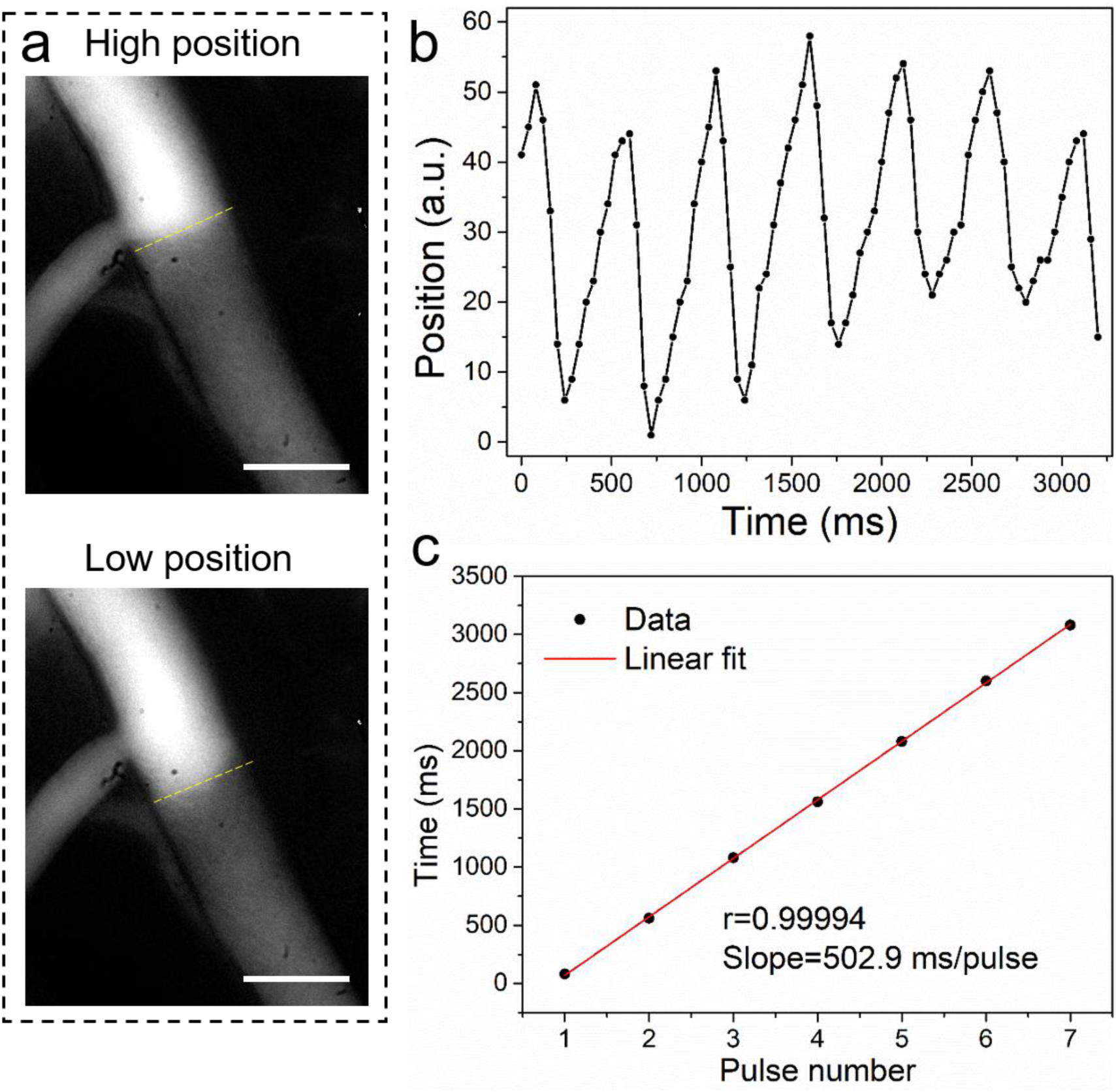
Measurement of cardiac impulse period of the rhesus macaque based on NIR-II fluorescence wide-field microscopic brain vascular imaging. **a** Two typical images with the fluorescence bright/dark border in the highest (top image) and lowest (bottom image) position in one cerebral blood vessel. Scale bars: 100 μm. **b** A plot of the position of the bright/dark boundary in the vessel as a function of time. **c** A plot of the peak timepoints of each impulse shown in **b**. The linear fit reveals an average cardiac impulse period of 502.9 ms/pulse, matching the 120 pulses/minute recorded on the heart rate monitor.

## NIR-II fluorescence confocal microscopy in mice

Prior to using the NIR-II fluorescence confocal microscope in primates, we first evaluated its performance in mice. As shown in Supplementary Fig. 6 and Supplementary Fig. 7, we imaged cerebral vasculature through a cranial window on a mouse which had received an intravenous injection of ICG (1 mg/mL, 200 μL). Images at various depths were obtained and a 3D image was reconstructed, revealing a clear vascular network (Supplementary Fig. 6). In addition, the spatial resolution and SBR were analyzed (Supplementary Fig. 7). This verified the superior imaging performance of our confocal setup.

**Supplementary Figure 6.**
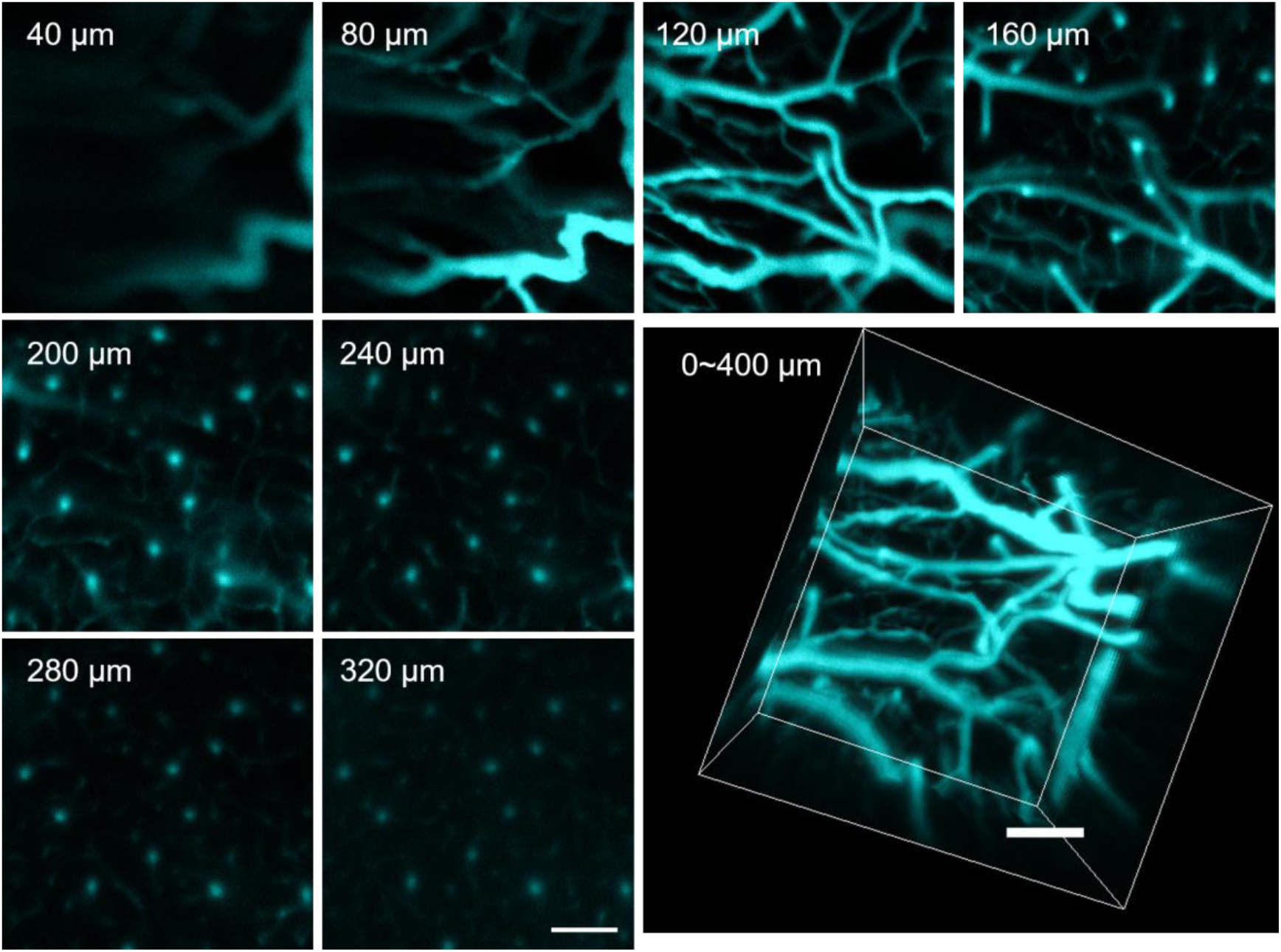
NIR-II fluorescence confocal microscopic images of cerebral blood vessels of mouse at various depths and a 3D reconstructed image over depths of 0-400 μm. Scale bars: 100 μm.

**Supplementary Figure 7.**
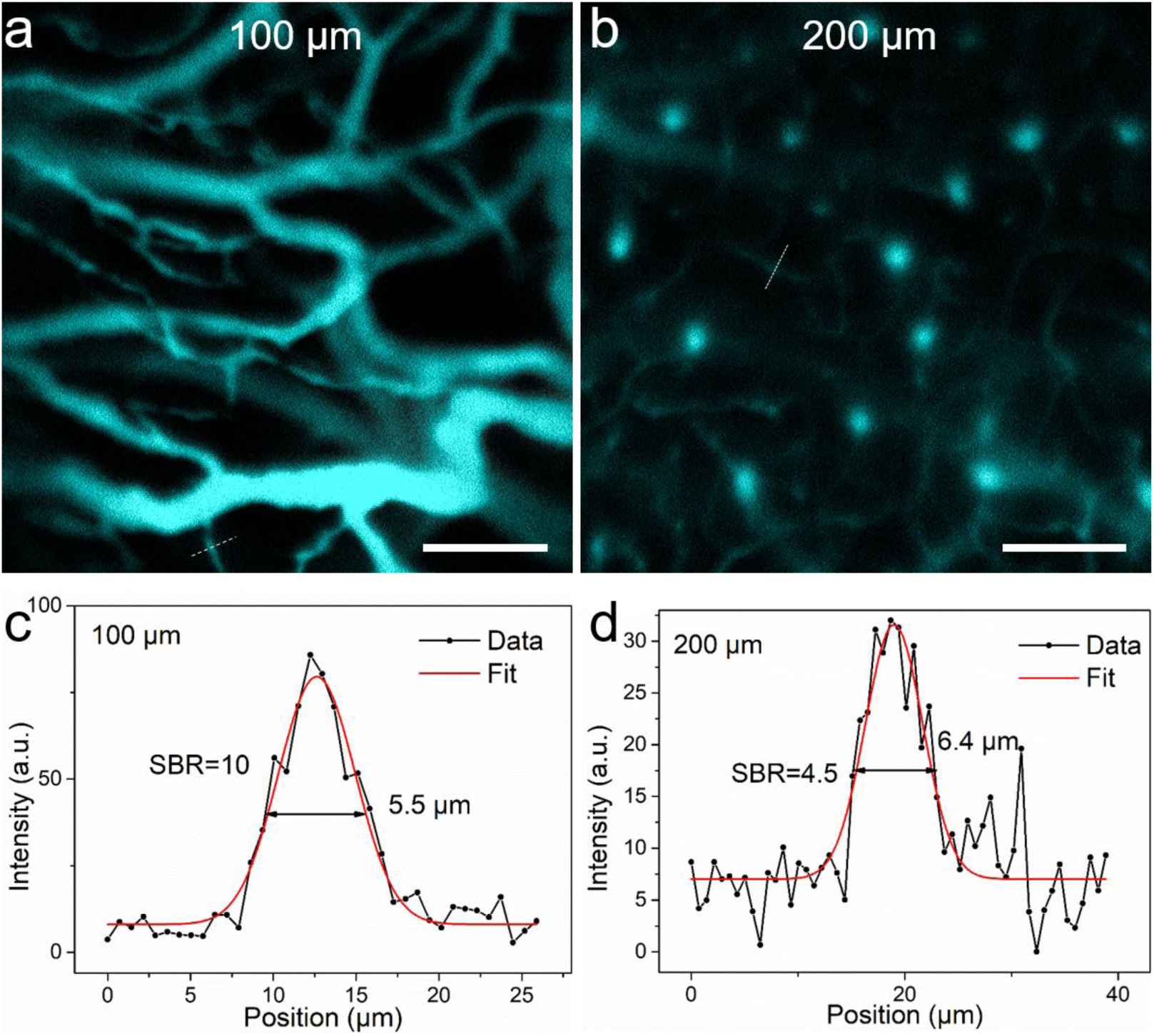
**a** and **b** NIR-II fluorescence confocal microscopic images of cerebral blood vessels of the mouse at two typical depths (100 μm and 200 μm). **c-d** The cross-sectional fluorescence intensity profiles (black) and the related Gaussian fitting (red) along the capillary vessels indicated by the white-dashed lines in **a** and **b**. Scale bars: 100 μm.

**MOV S1.** A movie showing the NIR-II fluorescence wide-field microscopic *in vivo* imaging the flow of cerebral blood vessels in the rhesus macaque, with the objective magnification of 25×.

**MOV S2.** NIR-II fluorescence wide-field microscopic *in vivo* imaging showing the blood flow directions in cerebral vessels, as well as the discrimination of artery and vein.

**MOV S3.** NIR-II fluorescence wide-field microscopic *in vivo* imaging showing the moving of fluorescence bright/dark border in one cerebral blood vessel acompanied with the cardiac impulse.

**MOV S4.** 3D reconstructed NIR-II fluorescence confocal microscopic *in vivo* images of cerebral blood vessels of the rhesus macaque.

